# Cooperative escape in ants and robots

**DOI:** 10.1101/2021.07.12.451633

**Authors:** S Ganga Prasath, Souvik Mandal, Fabio Giardina, Jordan Kennedy, Venkatesh N. Murthy, L Mahadevan

## Abstract

The solution of complex problems by the collective action of simple agents in both biologically evolved and synthetically engineered systems involves cooperative action. Understanding the resulting emergent solutions requires integrating across the organismal behaviors of many individuals. Here we investigate an ecologically relevant collective task in black carpenter ants *Camponotus pennsylvanicus*: escape from a soft, erodible confining corral. Individual ants show a transition from individual exploratory excavation at random locations to spatially localized collective exploitative excavation and escape from the corral. A minimal continuum theory that coarse-grains over individual actions and considers their integrated influence on the environment leads to the emergence of an effective phase space of behaviors in terms of excavation strength and cooperation intensity. To test the theory over the range of predicted behaviors, we used custom-built robots (RAnts) that respond to stimuli and show the emergence (and failure) of cooperative excavation and escape. Overall, our approach shows how the cooperative completion of tasks can arise from relatively simple rules that involve the interaction of simple agents with a dynamically changing environment that serves as an enabler and modulator of behavior.

---

Collective behavior in societies leads to the emergence of solutions to problems associated with brood care, foraging for food, protection from enemies and predation of prey, building complex architectures, and the cooperative resolution of physiological problems that are almost impossible to solve by individuals [1–6]. This benefits the society at large, and although there is a cost of cheating by non-participators, cooperation overall is known to play a critical role in the formation and sustenance of the social entities [1]. Cooperative behavior is seen across biological scales - from unicellular bacterium and slime molds, to animal societies and human organizations [7, 8]. Understanding the fundamental principles of cooperation and its breakdown [9, 10] links research in biology, mathematics, economics, policy-making and artificial intelligence, and takes many forms - from concrete studies that range from proximate genetic switches [11] for social behavior in ants to mechanisms underlying events like group hunting in wolves [12], transferring food by ants [13]; to cooperative brood care and division of labor in social groups [5, 14]; to abstractions [1] associated with the ultimate causes of the evolution of eusociality [15].

Social insects, and ants in particular, lend themselves to the study of cooperative problem solving, given their super-organismic colony structure, documented cooperative behavior, long evolutionary history and the relative ease of working with them in laboratory settings [16–18]. Minimally, task completion requires the ability of individuals to respond to local stimuli with stereotypical responses, communication between individuals that might involve using the environment, and a means to sense task execution and completion. For example, in the context of building complex architectures by termites, the presence (or absence) of pheromones serve as stimuli that communicate when and where, how to start (or stop) building [19, 20], while in ventilation and mechanoadaptation in bees, temperature, airflow and mechanical strain serve similar functions to provide collective solutions to the regulation of temperature, carbon dioxide or mechanical loads [3, 21, 22]. In all these cases, a dynamic and malleable environment is used as a communication channel, broadening the classical notion of stigmergy to include signaling via chemical, mechanical and fluidic means. But how individuals switch from local uncoordinated behavior to collective cooperation that translates to adaptable task execution in different social systems remains a relatively unexplored question.

Here we study an ecologically relevant task in carpenter ants *Camponotus pennsylvanicus*: escape from a confined corral via excavation and tunneling, using a combination of quantitative imaging in combination with mathematical and computational models, and then synthesize this behavior using custom-built robots that can respond to each other and the environment. Our study provides a phase diagram for the emergence of different classes of collective behavior and task completion using a set of local rules.

We start with ants drawn from a mature colony of *C. Pennsylvanicus* that consist of a queen, the sole egg layer, and the workers from three morphologically different castes - major, media and minor [23]. Though all ants perform different tasks like foraging, nest-keeping, brood care to a varied degree, during excavation, major ants, equipped with their large mandibles, generally take the lead role in excavating, while media and minor ants transport the debris outside their nest. Our experiments consist of a dozen worker ants from the same colony that are anesthetized (using *CO*_2_) and then brought into a confining corral made out of agarose flanked between two hard plastic sheets, without visible light to mimic their natural environment in a nest; infrared light was used to monitor the experiment using video (see SI Fig. 1(*a*)). Once the ants regain activity (due to the introduction of *O*_2_), they stay still for a while before moving. Observations show that they first exhibit wall-following followed by one of the ants initiating an exploratory excavation at a random location along the corral (ref SI Fig. S1). After an initial exploratory phase the ants switch to an exploitative strategy to excavate a tunnel at a specific location and eventually breakthrough the corral (seen through the sequence in Fig. 1(*b*)).

**FIG. 1.**
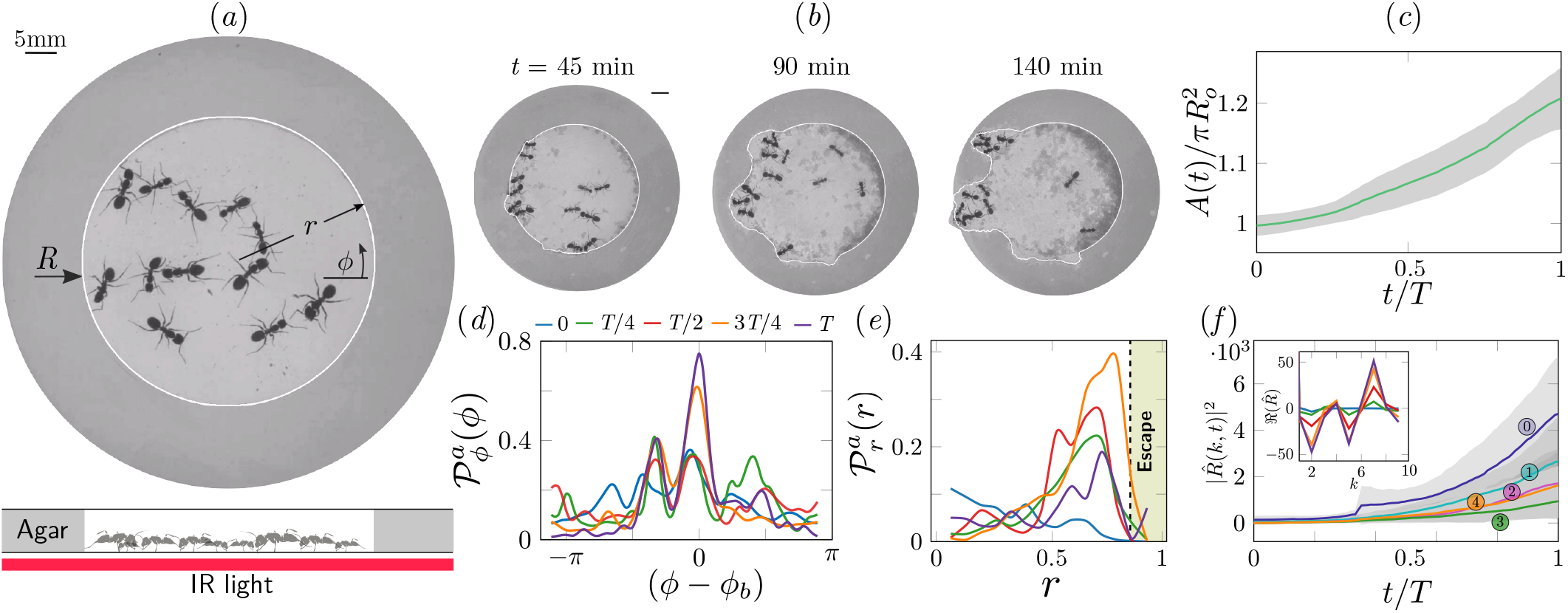
Collective dynamics of ant escape. (*a*) Colony members of the black carpenter ant *Camponotus pennsylvanicus* are confined to a porous boundary made out of Agarose. The boundary is represented by its radius *R*(*ϕ, t*) (*ϕ* - polar angle, *t* - time). Bottom part shows the side-view schematic of the experimental set-up with the boundary made of agarose and background IR light source used to image the ants in the dark. (*b*) Temporal progression of excavation experiments as 12 ants cooperatively tunnel through the agarose confinement. The white line is the tracked location of the inner wall which grows in size as the excavation progresses. (*c*) Confinement area *A*(*t*) as a function of time (scaled by escape time *T*), normalized by initial circular confinement with radius *R*_*o*_. (*d*) Evolution of the orientation distribution of the ant density, 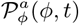 obtained by averaging along the radial direction. Ants start from an initially isotropic state and localize at an angle *ϕ*_*b*_ along the boundary. *T* here is the escape time. (*e*) Dynamics of the radial distribution of ant density 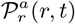 as a function of radial distance, *r* obtained by averaging a sector of π/6 around the excavation site. We see that the ant density front propagates through the corral to escape. The density is plotted for the same times as in (*d*). (*f*) Evolution of the power spectrum 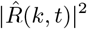 of first five Fourier modes capturing the number of tunnels formed during excavation 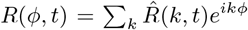. Inset shows the real part of the Fourier coefficient, 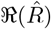 at different time instants indicating that many modes are present in the boundary shape.

We quantify this transition from rotationally isotropic exploration to localized excavation by considering the density of ants ϱ_*a*_(*r, ϕ, t*) as a function of space and time in the corral, using cylindrical coordinates (*r, ϕ*). Over time, the density becomes localized at a particular angle and location along with corral where large-scale excavation eventually leads to escape (see Video 3, SI Fig. S1 to get a coarse-grained view of the spatio-temporal evolution of the ant density, obtained by averaging over individual ant movements). Simultaneously, we see a signature of collective excavation in an increase of the volume of excavated material, as shown in Fig. 1(*c*). Averaging the density over radial positions, in Fig. 1(*d*) we show the orientation distribution of the ant density 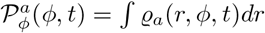 is initially isotropic, and gradually starts to localize at a particular (arbitrary) value of the angle as time increases.

Averaging the density over the localized region, in Fig. 1(*e*) we show the radial distribution of the ant density 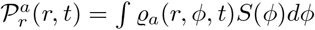 (where *S*(*ϕ*) is a kernel around the excavation site) that is initially uniform, and gradually propagates inside the boundary of the corral as time increases. Consistent with localization and concomitant excavation (Fig. 1(*f*) inset, SI Fig. S2(*c*)), we see that the Fourier amplitudes of multiple modes compete with each other initially before an elliptic mode (corresponding to a strongly localized state) is amplified as excavation progresses (shown in Figs. 1(*f*), SI Fig. S2(*b*)).

All together, our quantitative observations show that an initially isotropic and homogeneous distribution of ants in the corral induces exploration of multiple potential tunneling paths that transitions into the exploitative excavation of one specific location that eventually leads to an escape route. To understand this transition, we now turn to a minimal theoretical framework that coarse-grains over the fast times and short length scales (of ant motions and sizes). Our aim here is to show that an effective theory that does not dwell on the details of individual agents (ants) or their interactions with each other and the environment, but instead couples three slowly-varying fields: the ant density, a pheromone/antennating field that the ants use to communicate with each other, and the corral, suffices to explain our observations in terms of a small number of effective parameters.

We define these interacting spatio-temporal fields as ant density ϱ_*a*_(**x**, *t*), antennating field *c*(**x**, *t*) and corral density ϱ_*s*_(**x**, *t*) shown schematically in Fig. 2(*a*). In the absence of any external gradients, we assume that ants move randomly, but this diffusive movement is rectified by pheromone gradients or reinforcing antennating signals [16, 24–26], in addition to being self-propelled with a velocity **u**_*a*_ that is related to the local environment. Though antennation and pheromone are two different mechanisms of communication as the information is carried with the animal in the former and is left in the environment in the latter, their dynamics is expected to be same when the ants move slower than the time scale associated with memory diffusion or decay. In turn, pheromone signals are laid down by ants at a rate proportional to their density, and then degrade and diffuse slowly. Finally, the ant collective excavates the corral at a rate proportional to the difference in the pheromone concentration relative to a threshold value (see SI sec. S2 SS1). Accounting for these effects, we arrive at the following dynamical equations for the evolution of ϱ_*a*_(**x**, *t*), *c*(**x**, *t*) and ϱ_*s*_(**x**, *t*):

**FIG. 2.**
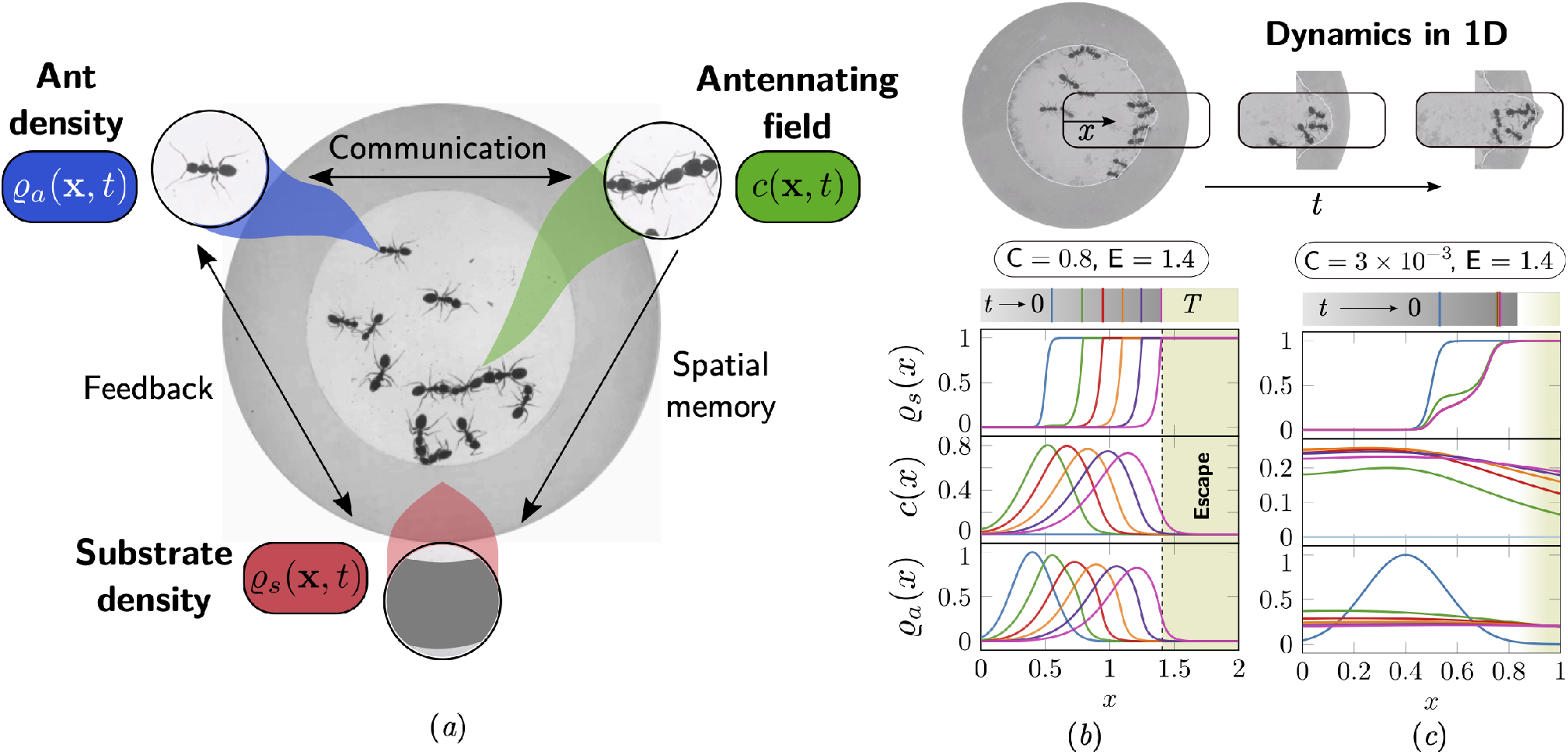
Cooperation via organism-environment-organism interaction. (*a*) Schematic of the model showing the interaction between the different spatio-temporal fields required to capture cooperative escape of ants: ant density, ϱ_*a*_(**x**, t); concentration of antennating field, *c*(**x**, t) capturing inter-ant communication; density of corral, ϱ_*s*_(**x**, t) representing the soft corral which the ants excavate. We capture the dynamics of excavation by ants close to the excavation site using the 1-dimensional version of Eqns. 1-3. (*b, c*) Temporal progression of the corral density, antennating field and the ant density showing successful escape for high cooperation captured using the non-dimensional number, C (representing non-dimensional strength of cooperation amongst ants) and faster excavation, captured using E. For reduced cooperation ants’ diffusion dominates and only partial tunnels are formed (see SI S3 for details). *T* here is the time of escape.

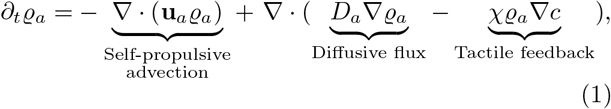

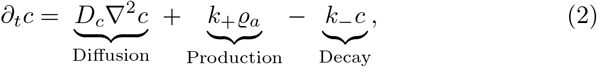

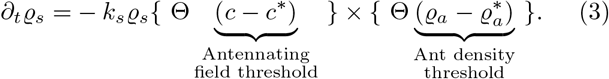

Here the ant advection velocity is assumed to have the form 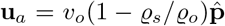 where *v*_0_ is the characteristic velocity of ants, and 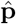 is a unit vector pointing along the *θ* direction, and the term (1 − ϱ_*s*_/ ϱ_*o*_) reflects the fact that excavating ants are slowed down by their labor; *D*_*a*_ is the diffusivity of the ant, χ is the strength of antennating field following behavior; *k*_+_, *k*_−_ are the rate of production and decay of the antennating/pheromone field, and *D*_*c*_ is the diffusivity of the pheromone (see SI sec. S3 SS3); *k*_*s*_ is the rate of excavation of the corral and 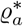, *c*^*^ are the threshold concentration of ant density and antennating field required to initiate excavation, that serve to trigger simple switch-like behaviors via the Heaviside function Θ(*x*) (or its regularization via hyperbolic or Hill functions). In the absence of excavation dynamics, our framework reduces to the well known Keller-Segel model for chemotaxis [26] (detailed in SI sec. S3). Coupling this to the dynamics of excavation introduces the all-important notion of *functional* collective behavior linking active agents, communication channels (the antennating and pheromone fields) and a dynamic, erodible corral that characterizes function in terms of progress towards task completion.

Although our mdoel has a number of dimensionless parameters (see SI sec. S3 for a list and ranges), just two are qualitatively important in capturing the ethology of cooperative excavation: (i) the scaled cooperation parameter defined as C = χ*c*_*o*_/*D*_*a*_ which determines the relative strength of antennation (gradient-following) to ant diffusion, (ii) the scaled excavation rate, E = *k*_*s*_*l*/*v*_*o*_. Here, *l*/*v*_*o*_ is the characteristic time-scale of ant motion, with *l* ∼ min[(*D*_*c*_/*k* _)^1*/*2^, *l*_*a*_], where *l*_*a*_ is the ant size (see SI sec. S3 for details). Solving the governing Eqns. 1-3 in a one-dimensional setting (ref SI sec. S3 SS4) captures the two limits of the excavation behavior and escape seen in experiments; for large excavation rate E and cooperation parameter, C, we see successful escape (shown in Fig. 2(*b*)), while decreasing the cooperation parameter leads to a failed strategy (shown in Fig. 2(*c*)). All together, our phase-field model shows the emergence of cooperativity without the need for a plan, optimization principle, or an internal representations of the world, but via the environmentally-mediated communication between agents [27] that leads to task completion.

To go beyond our ability to explain the observations of ant behavior using our theoretical framework, we need to probe a larger range of the parameters and phase-space spanned by C, E, than our experiments allowed us to. For this, we turn to a robotic platform to synthesize collective functional behaviors that arise from simple behavioral rules underlying individual programmable robots. Our custom designed robot ants (RAnts) are inspired by many earlier attempts to create artificial agents that are mobile and follow simple rules [28–30], can respond to virtual pheromone fields [31, 32] and are capable of robotic excavation [33]. Our autonomous wheeled robots can exhibit emergent embodied behavior [34, 35], and are flexible enough to allow for a range of stigmergic interactions with the environment [36, 37]. This is made possible by having each RAnt equipped with an infrared distance sensor to detect obstacles and other RAnts, a retractable magnet that can pick up and drop wall elements with a ferromagnetic ring (shown in Fig. 3(*a*)), and the ability to measure a virtual pheromone field generated by a light projected (from below) onto the surface of a transparent arena they operate in (see Fig. 3(*a, b*)) [31, 32, 38, 39]. The intensity of this “photormone” field follows the antennating field Eq. 2. This allows us to use a local form of Eqns. 1-3 to define a robot’s behavior in terms of an excavation rate E, a cooperation parameter C, and a threshold concentration for tunneling *c*^*^, encoded in terms of the following behavioral rules: (i) follow gradient of projected photormone field; (ii) avoid obstacles and other RAnts at higher photormone locations; (iii) pick up obstacles from high photormone locations and drop them at low concentration levels (see Fig. 3(*c*) and SI sec. S4 for more details).

**FIG. 3.**
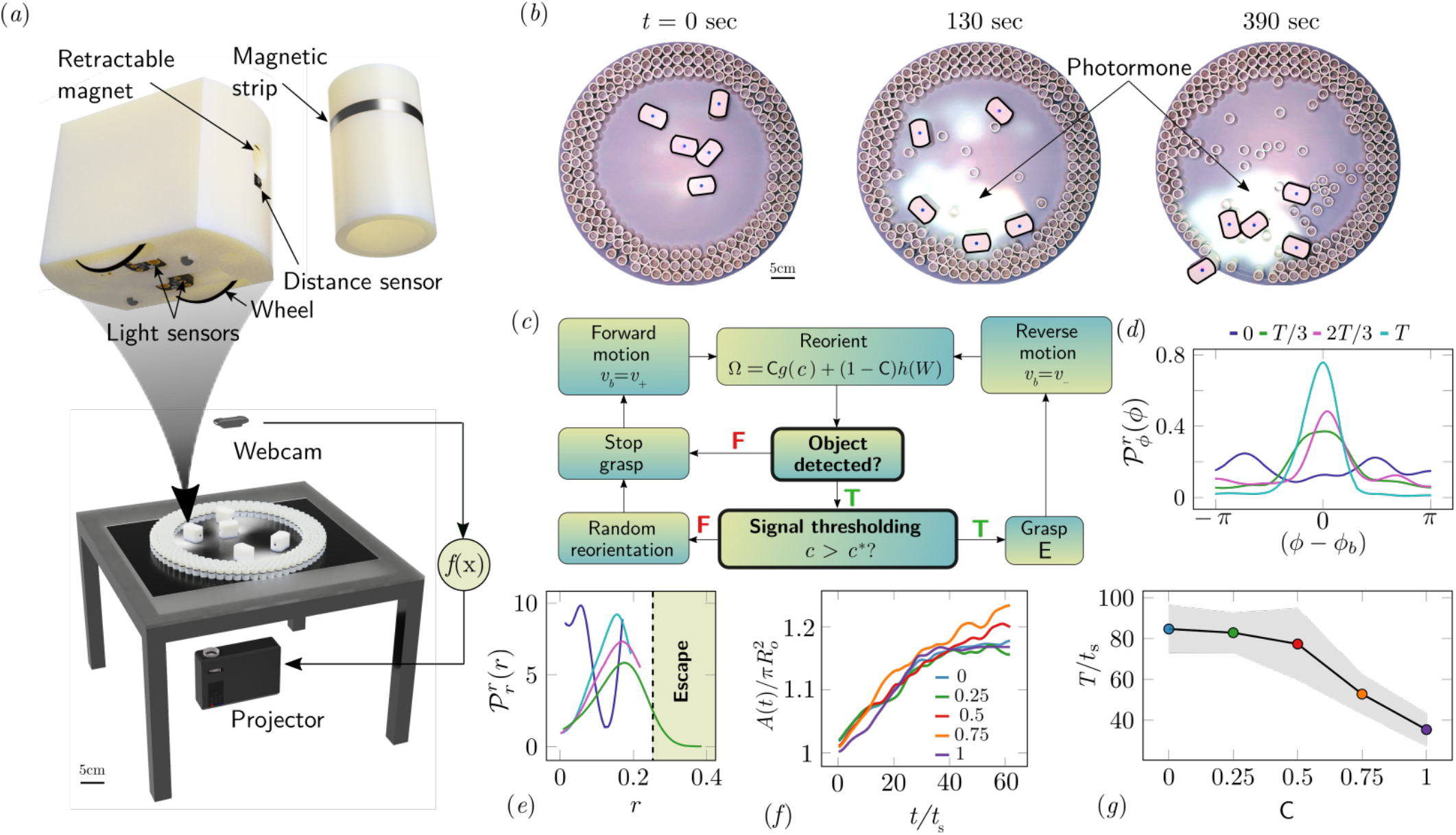
Emergent escape dynamics in robotic ants. (*a*) Robot Ant (*RAnt*) set-up. A mobile RAnt is placed in an arena 50cm in diameter surrounded by three layers of cylindrical boundary elements totalling 200 elements. The outermost layer is prevented from being pushed out of the arena by a circular ring. A scalar concentration field (*photormone* field) is projected onto a plane whose intensity can be measured by a RAnt. The position of each RAnt is tracked using a webcam. Each RAnt can pick up and drop the discrete boundary elements using a retractable magnet. (*b*) Series of snapshots at different times of the escape process for a cooperation parameter C = 1. (*c*) Flowchart of the RAnt programming. A base locomotion speed v_*b*_ is stored internally and the rate of change Ω of the heading is a function of the cooperation parameter C, the photormone concentration *c*, and a stochastic process *W* (Brownian motion). A photormone threshold *c*^*^ determines whether an object is grasped (with probability E) after it is detected by the distance sensor. (*d*) Orientation distribution of the RAnt density 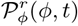 as a function of the azimuthal position *ϕ. ϕ*_*b*_ is the orientation of the escape tunnel. The density is plotted for different times. (*e*) Radial distribution of the RAnt density 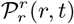 within a sector of π/2 centered around the position of the escape tunnel as a function of distance from the center of the arena *r*. The density is plotted for the same times as in (*d*). (*f*) Confinement area *A*(*t*) as a function of time, normalized by initial circular confinement with radius *R*_*o*_ for different cooperation parameter C. (*g*) Normalized escape time *T* as a function of cooperation parameter C, averaged over 5 experiments per cooperation parameter. Every experiment was run until the first RAnt escaped or the experiment duration exceeded 15 minutes.

Varying the parameter C ∈ [0, 1] allows us to dial the individual behavior from random motion (C = 0) to tracking the photormone gradient (C = 1), while varying the non-dimensional excavation rate E by changing the frequency at which the robots execute pick-and-drop behavior with the boundary wall serves to mimic what arises in ants as a function of their morphology and caste (see SI sec. S2 SS1 for more details). For a specific value of these parameters, we followed the collective behavior of RAnts by averaging their position over several pick-and-drop timescales to obtain the RAnt density field ϱ_*r*_(*r, ϕ, t*), just as for ants.

When all the RAnts are programmed to have a cooperation parameter C = 1, RAnts initially explore the region without picking the boundary element until the photormone concentration c ∼ c^*^, which happens once a particular location has enough visits by other RAnts. As for ants, we calculate the radially averaged RAnt density 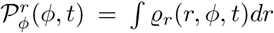; Fig. 3(d) shows how RAnt density localizes at a (random) value of the azimuthal angle. As excavation progresses, the RAnt density propagates radially outwards as a density front just as in ants, shown in Fig. 3(e) in terms of the quantity 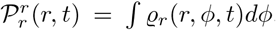. Concommitantly, as excavation progresses, the corral area increases; interestingly the scaled corral area 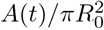 is independent of the cooperation parameter C as shown in Fig. 3(*f*) (all RAnts were programmed to have the same excavation rate). However, cooperation does change the time for escape; in Fig. 3(*g*) we show the average escape time (scaled by the characteristic time it takes for a rant to traverse the arena) and see that *T*/*t*_s_ decreases with an increase in the cooperation parameter C. RAnts escaped every time for C > 0.5, but are unable to escape for low cooperation parameters (within a 15 minute time window). Our results show that it is the localized collective excavation of RAnts mediated by photormone-induced cooperation that is responsible for efficient tunneling and escape; for low values of the cooperation parameter, tunneling is defocused and global, and thus not as effective (see Fig. S7). When E → 0 (vanishing probability for a successful pick up) but strong cooperation (see Fig. 4 and SI sec. S3 SS3 for theoretical predictions), the RAnts get jammed because they follow the photormone field they generate but are unable to tunnel through the boundary constriction. On the other hand, when E is small and C is small, the agents do not cooperate and their diffusive behavior prevents successful tunneling. The range of strategies can be visualized in a two-dimensional phase space spanned by the variables E and C shown in Fig. 4. Low values of C and E lead to diffusive (and non-functional) behavior, while high values of these variables lead to successful escape, with the other two quadrants corresponding to jammed states (large C, small E) and partially tunneled states (large E, small C). These last two states are also observed as transients in ant collectives (see Video3).

**FIG. 4.**
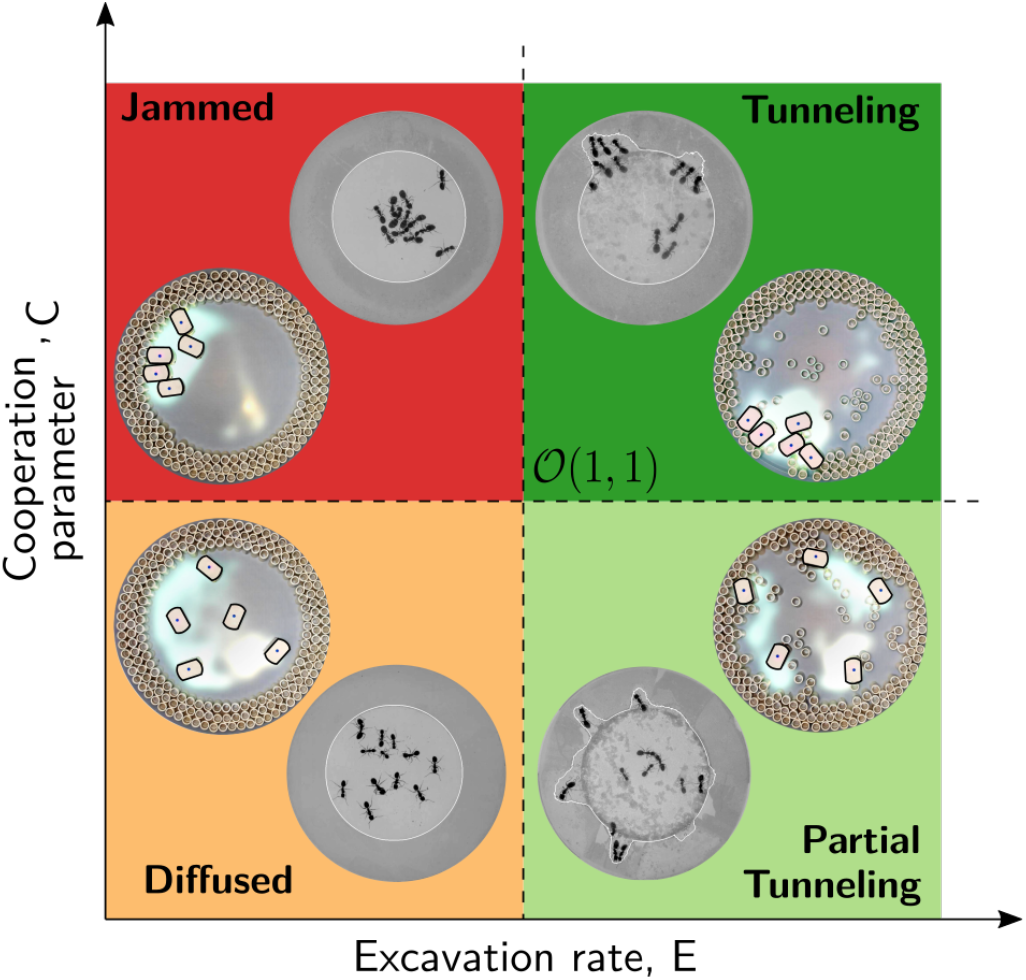
Phases of cooperation. Phase-diagram of cooperative task execution with different phases seen in ants and RAnts. In the robotic experiments we tune the Cooperation parameter C and the Excavation rate E while in the ant experiments we change the caste mixture. In the ant experiments we see the jammed and diffused phases transiently before the ants relax to cooperative excavation.

More than half a century ago, Tinbergen laid down a few basic principles for studying behavior [40] that focus on its function, mechanism, development and evolution. To this, we may add one more: synthesis. This naturally requires an elucidation of the underlying mathematical, physical and engineering principles that provide minimal mechanisms and thus suggest pathways for the development and evolution of complex behaviors, and create their robotic mimics. In the current context, the excavating behavior of individual ants serves to facilitate the function of escape from confinement, but becomes viable only through the collective action of a group. Our observations show how the transition from an individually exploratory to an exploitative cooperative tunneling strategy is mediated by the local chemical and mechanical environment. A simple dynamical model explains our observations and provides a minimal phase diagram for a range of strategies. To test and generalize this, we synthesized the observed behavior using robotic mimics that follow a minimal set of behavioral rules that mould the environment and are modulated by it. Our theoretical and robotic framework relies on simple local rules and a malleable environment that serves as both part of the system memory as well as a computational platform (using the spatio-temporal photormone field and the corral). It is also robust to failure of and stochasticity in the behavior of individual agents, in the communication channels or in the corral geometry, and instead considers only coarse-grained variables, in sharp contrast to engineering approaches that aim to control all agents and optimize costs. Different strategies such as escape, jamming and diffusion then arise as a function of the relative strength of the cooperation (representing the ability to follow/remember gradients) and excavation parameters (representing the ability to move material), as manifest in a simple phase diagram, and the emergence of cooperation arises due to the relatively slow decay of an environmental signal (the pheromone/antennating/photormone field), coupled to a threshold excavation rate. Our approach to functional and purposeful collective behavior links many simple brains and bodies with a dynamic environment that modulates behavior, and is changed by it. Since the ability to solve complex eco-physiological problems such as collective escape is directly correlated with a selective advantage in an evolutionary setting, perhaps collective behavior must always be studied in a functional context.

## ACKNOWLEDGMENTS

We thank the NSF PHY1606895 (S.G.P., L.M.), Swiss National Science foundation (F.G., grant P400P2-191115), Ford foundation (J.K.), NSF EFRI 18-30901 (L.M.), NSF 1764269 (L.M.), Kavli Institute for Bionano Science and Technology (S.M., V.M., L.M.), and the Henri Seydoux Fund (L.M.) for partial financial support.

## References

[1] M. A. Nowak, Evolutionary dynamics: exploring the equations of life (Harvard university press, 2006).

[2] O. Feinerman, I. Pinkoviezky, A. Gelblum, E. Fonio, and N. S. Gov, Nature Physics 14, 683 (2018).

[3] S. A. Ocko and L. Mahadevan, Journal of The Royal Society Interface 11, 20131033 (2014).

[4] S. Camazine, J.-L. Deneubourg, N. R. Franks, J. Sneyd, G. Theraulaz, and E. Bonabeau, Self-organization in biological systems (Princeton university press, 2020).

[5] B. Hölldobler, E. O. Wilson, et al., The superorganism: the beauty, elegance, and strangeness of insect societies (WW Norton & Company, 2009).

[6] H. King, S. Ocko, and L. Mahadevan, Proceedings of the National Academy of Sciences 112, 11589 (2015).

[7] J. Alcock, Animal behavior: An evolutionary approach (Sinauer Associates Sunderland, 2001).

[8] E. Pennisi, Science 325, 1196 (2009)

[9] J. Elster et al., Social mechanisms: An analytical approach to social theory (Cambridge University Press, 1998).

[10] I. D. Couzin, J. Krause, et al., Advances in the Study of Behavior 32, 10 (2003).

[11] W. Trible, L. Olivos-Cisneros, S. K. McKenzie, J. Saragosti, N.-C. Chang, B. J. Matthews, P. R. Oxley, and D. J. Kronauer, Cell 170, 727 (2017).

[12] C. Muro, R. Escobedo, L. Spector, and R. Coppinger, Behavioural processes 88, 192 (2011).

[13] H. F. McCreery and M. Breed, Insectes sociaux 61, 99 (2014).

[14] A. Brahma, S. Mandal, and R. Gadagkar, Proceedings of the National Academy of Sciences 115, 756 (2018).

[15] M. A. Nowak, C. E. Tarnita, and E. O. Wilson, Nature 466, 1057 (2010).

[16] B. Hölldobler, E. O. Wilson, et al., The ants (Harvard University Press, 1990).

[17] D. M. Gordon, Ants at work: how an insect society is organized (Simon and Schuster, 1999).

[18] A. Perna and G. Theraulaz, Journal of Experimental Biology 220, 83 (2017).

[19] S. A. Ocko, A. Heyde, and L. Mahadevan, Proceedings of the National Academy of Sciences 116, 3379 (2019).

[20] A. Heyde, L. Guo, C. Jost, G. Theraulaz, and L. Mahadevan, Proceedings of the National Academy of Sciences 118, e2006985118 (2021).

[21] O. Peleg, J. M. Peters, M. K. Salcedo, and L. Mahadevan, Nature Physics 14, 1193 (2018).

[22] J. M. Peters, O. Peleg, and L. Mahadevan, Journal of the Royal Society Interface 16, 20180561 (2019).

[23] L. D. Hansen and J. H. Klotz, Carpenter ants of the United States and Canada (Cornell University Press, 2005).

[24] J. Reinhard and M. V. Srinivasan, Food exploitation by social insects: ecological, behavioral, and theoretical approaches 1, 165 (2009).

[25] C. M. Waters and B. L. Bassler, Annu. Rev. Cell Dev. Biol. 21, 319 (2005).

[26] T. Hillen and K. J. Painter, Journal of mathematical biology 58, 183 (2009).

[27] M. J. Mataric, in Proceedings of the Second International Conference on Simulation of Adaptive Behavior (1993) pp. 432–441.

[28] V. Braitenberg, Vehicles: Experiments in synthetic psychology (MIT press, 1986).

[29] R. A. Brooks, Artificial intelligence 47, 139 (1991).

[30] H. A. Simon, The sciences of the artificial, 3rd ed. (MIT press, 1996).

[31] K. Sugawara, T. Kazama, and T. Watanabe, in International Conference on Intelligent Robots and Systems, Vol. 3 (IEEE, 2004).

[32] S. Garnier, F. Tache, M. Combe, A. Grimal, and G. Theraulaz, in Swarm intelligence symposium (IEEE, 2007).

[33] J. Aguilar, D. Monaenkova, V. Linevich, W. Savoie, B. Dutta, H.-S. Kuan, M. Betterton, M. Goodisman, and D. Goldman, Science 361, 672 (2018).

[34] L. Giomi, N. Hawley-Weld, and L. Mahadevan, Proceedings of the Royal Society A: Mathematical, Physical and Engineering Sciences 469, 20120637 (2013).

[35] A. Bricard, J.-B. Caussin, N. Desreumaux, O. Dauchot, and D. Bartolo, Nature 503, 95 (2013).

[36] J. Werfel, K. Petersen, and R. Nagpal, Science 343, 754 (2014).

[37] K. H. Petersen, N. Napp, R. Stuart-Smith, D. Rus, and M. Kovac, Science Robotics 4 (2019).

[38] G. Theraulaz and E. Bonabeau, Science 269, 686 (1995).

[39] G. Wang, T. V. Phan, S. Li, M. Wombacher, J. Qu, Y. Peng, G. Chen, D. I. Goldman, S. A. Levin, R. H. Austin, et al., Physical review letters 126, 108002 (2021).

[40] N. Tinbergen, The study of instinct (Pygmalion Press, an imprint of Plunkett Lake Press, 2020).

[41] T. D. Pereira, N. Tabris, J. Li, S. Ravindranath, E. S. Papadoyannis, Z. Y. Wang, D. M. Turner, G. McKenzie-Smith, S. D. Kocher, A. L. Falkner, et al., bioRxiv (2020).

[42] M. Le Goc, L. H. Kim, A. Parsaei, J.-D. Fekete, P. Dragicevic, and S. Follmer, in Proceedings of the 29th Annual Symposium on User Interface Software and Technology (2016) pp. 97–109.

